# Differentiation Protocol-Dependent Variability in hiPSC-Derived Endothelial Progenitor Functionality

**DOI:** 10.1101/2024.09.27.615423

**Authors:** Brett Stern, Sarah Meng, Bryce Larsen, Amy Brock, Janet Zoldan

**Affiliations:** Department of Biomedical Engineering, University of Texas at Austin, Austin, Texas, USA

## Abstract

**Purpose:** With multiple groups generating methods for differentiating induced pluripotent stem cells (hiPSCs) into endothelial cells/endothelial progenitors (ECs/EPs), there is a need to better understand the specific endothelial subtypes that different differentiation protocols produce. This is especially important for accurate tissue modeling as researchers continue to incorporate the endothelium into engineered tissues.

**Methods:** We illustrated the heterogeneity of cells produced from different differentiation protocols by focusing on two selected protocols, one driving differentiation exclusively using small molecules (SM Protocol) and one driving differentiation primarily using growth factors (GF Protocol). We characterized the cells through a combination of vasculogenic computational analysis following encapsulation in 3D hydrogels, RNA sequencing, and measurement of soluble factor release.

**Results:** Vasculogenic computational analysis indicates that cells from the GF protocol formed more dense and interconnected vasculature compared to cells from the SM protocol. Likewise, RNA-seq analysis showed GF-derived cell enrichment in pathways involved in cell migration and angiogenesis. In addition, GF-derived cells favor differentiation to arterial endothelial cells as well as predisposition to undergoing a partial endothelial-to-mesenchymal transition, whereas SM-derived cells resemble immature progenitors based on higher proliferation rates as well as ECM remodeling. These trends also persisted following extended 3D culture.

**Conclusions:** The results demonstrate that despite using the same starting hiPSC population and isolating EPs using the same surface marker, the two differentiation protocols yield highly distinct cell populations. This work highlights the importance of understanding the specific endothelial subtypes produced by different differentiation protocols for the creation of more accurate tissue models.

**Lay Summary:** Although multiple groups have created protocols for differentiating human induced pluripotent stem cells (hiPSCs) to endothelial progenitors (EPs), it is not clear if different differentiation protocols generate different endothelial subtypes, matching the variance seen *in vivo* To that end, we performed an in-depth characterization of two selected hiPSC-EP differentiation protocols, primarily focusing on cells cultured in 3D hydrogels, to better understand the specific endothelial subtypes produced. We found significant differences in both cell maturity and cell phenotype that persisted even after extended culture.

**Future Work:** Future work will involve extending the existing study to include additional hiPSC-EP differentiation protocols as well as conducting single-cell RNAseq to better understand the makeup of different cell populations both before and after isolating CD34^+^ cells. Because smooth muscle cells play an important role in vessel stabilization, we will also incorporate hiPSC-derived smooth muscle cells into the system.

## Introduction

Human induced pluripotent stem cells (hiPSCs) show considerable promise for numerous applications, ranging from disease modeling and drug discovery to regenerative medicine. For vascular applications, hiPSCs can be differentiated into endothelial cells (ECs), which form perfusable vasculature both *in vitro* and *in vivo* Multiple groups have developed protocols for differentiating hiPSCs to ECs through small molecule, growth factor, and even plasmid supplementation^1-12^ and verified their functionality through a variety of methods.

One key challenge in using hiPSC-ECs for tissue engineering is the wide heterogeneity of endothelial cell populations *in vivo* This includes both arterial/venous specification for larger diameter vessels as well as tissue specific phenotypes associated with capillaries. Although all endothelial cells express common surface markers^13^, such as CD34, PECAM, and VE-Cadherin, expression levels vary between ECs in different tissues. These differences also extend to the morphological level, with three major classifications of vessels: continuous, fenestrated, and discontinuous. As the field advances and generates more complex tissue structures, there is a greater need to incorporate tissue-specific endothelial cells.

We can leverage existing hiPSC-EC differentiation protocols to address this issue. Several groups have focused on generating tissue-specific ECs^13^, mostly focusing on arterial/venous specification, through modifications to various aspects of the differentiation protocol, such as small molecules and/or growth factors used to drive differentiation, length of time for differentiation, and surface marker(s) for identifying successfully differentiated cells. However, there are a significant number of differentiation protocols, including those with widespread adoption, in which the cell types produced are not well established. As a result, it is likely that different differentiation protocols are producing significantly different endothelial populations.

To illustrate the heterogeneity of hiPSC-EP populations, we conducted a thorough characterization of two selected hiPSC-EP differentiation protocols that differ only in the factors used for differentiation: one drives differentiation through small molecules (SM protocol) and the other drives differentiation through growth factors (GF protocol). We found that, despite using the same starting hiPSC line, differentiating over the same length of time, and sorting using the same surface marker, these two differentiation protocols yield vastly different cell populations. Specifically, we found that cells derived using the GF protocol form more robust vasculature when encapsulated in 3D hydrogel scaffolds and are predisposed to undergoing an endothelial-to-mesenchymal transition, whereas cells derived using the SM protocol are more immature and stem-like. These trends also persist following extended culture. More broadly, this work demonstrates the need for more robust characterization of both preexisting and newly developed hiPSC-EP differentiation protocols to better understand the specific endothelial subtypes produced; by extension, researchers using hiPSC-derived EPs should also more carefully consider their choice of differentiation protocol and tailor it to their specific applications.

## Materials and Methods

### Maintenance of hiPSCs

hiPSCs (WiCell, DF19-9-11T) were cultured on vitronectin-coated (ThermoFisher Scientific, A14700) six-well plates in Complete Essential 8 medium (E8, A1517001, ThermoFisher Scientific). hiPSCs were passaged upon reaching 70–80% confluency. To passage, hiPSCs were treated with 0.5 mM ethylenediaminetetraacetic acid (EDTA, Invitrogen, 11140-036) for 4 minutes at 37°C. EDTA was then removed and the hiPSCs were resuspended in Complete E8 and seeded as small colonies onto fresh vitronectin-coated plates.

### Differentiation of hiPSCs to CD34^+^-hiPSC-EPs Using Small Molecules (SM Protocol)

hiPSCs were differentiated into hiPSC-EPs following a modified protocol from Lian *et al*^12^. hiPSCs were manually dissociated into a single-cell suspension in Complete E8 supplemented with 10 µM Y-27632 (Selleckchem, S1049) and plated on Matrigel-coated (Corning, 356231) 6-well plates at a density of 20,000 cells/cm^2^. 24 hours after seeding, media was replaced with Complete E8 without Y-27632. 48 hours after seeding, media was replaced with LaSR Basal and 6 µM CHIR99021 (LC Laboratories, C-6556). LaSR Basal is comprised of Advanced Dulbecco*’*s Modified Eagle Medium (DMEM)/F12 (ThermoFisher Scientific, 12634010) supplemented with 60 µg/mL L-ascorbic acid 2-phosphate (Sigma-Aldrich, A8960) and 2.5 mM GlutaMAX (ThermoFisher Scientific, 35050061). 48 hours after CHIR99021 induction, media was replaced with LaSR Basal without CHIR99021, and the cells were cultured for three additional days.

### Differentiation of hiPSCs to CD34^+^-hiPSC-EPs Using Growth Factors (GF Protocol)

hiPSCs were differentiated into CD34^+^-hiPSC-EPs using a modified protocol from Jalilian *et al*^1^. To summarize, hiPSCs were dissociated into a single cell suspension and plated onto Matrigel-coated 6-well plates at a density of 10,000 cells/cm^2^ in Complete E8 supplemented with 10 µM Y-27632. 24 hours after seeding, media was replaced with Complete E8 without Y-27632. 48 hours after seeding, media was replaced with Differentiation Media supplemented with 25 ng/mL Activin A (R&D Systems, 330-AC-050), 30 ng/mL Bone Morphogenic Protein 4 (BMP4, R&D Systems, 314-BP-050), and 0.15 µM 6-bromoindirubin-3-oxime (BIO, Selleckchem, S7198). Differentiation Media consists of Dulbecco*’*s Modified Eagle Medium (DMEM)/F12 (Cytiva, SH30023.FS) supplemented with N2 supplement (ThermoFisher Scientific, 17502048) and B27 Supplement minus Insulin (ThermoFisher Scientific, A1895601). Differentiation Media supplemented with Activin A, BMP4, and BIO was replaced after 24 hours. 96 hours after seeding, media was replaced with Differentiation Media supplemented with 2 µM SB431542 (Tocris, 1614) and 50 ng/mL Vascular Endothelial Growth Factor (VEGF, R&D Systems, 203-VE-050). Differentiation Media supplemented with these factors was replaced daily for 2 additional days.

### Fluorescence-activated cell sorting (FACS) of CD34^+^-hiPSC-EPs

7 days after seeding, the differentiated cells were incubated with Accutase (STEMCELL Technologies, 07920) for 10 minutes at 37°C and dissociated into single cells. Cells were centrifuged at 300g for 5 minutes and the pellet was resuspended in 200 µL of sorting buffer, containing 2 mM EDTA and 0.5% bovine serum albumin (BSA; Sigma-Aldrich, A8412-100ML) in Dulbecco*’*s Phosphate-Buffered Saline (DPBS, GE Healthcare, SH30028.03), and 2 µL of CD34-PE antibody (Miltenyi Biotec, 130-113-741). Cells were then incubated at 4°C for 10 minutes. Cells were filtered twice with a 35 µm cell strainer (Corning, 352235) to remove cell and extracellular matrix aggregates. CD34^+^-hiPSC-EPs were isolated using a fluorescence-activated cell sorting instrument (S3e; Bio-Rad). Gating and population analysis were performed with Bio-Rad software native to the S3e cell sorter.

### Optimization of Growth Factor Differentiation Protocol

In order to determine the factors most important to maximize CD34^+^-hiPSC-EP yield, we used Minitab Statistical Software to perform a screening test. We tested a total of seven factors: seeding density (10,000 or 40,000 cells/cm^2^), B-27 supplement (with or without insulin), Activin A (0 or 25 ng/mL), BMP4 (0 or 30 ng/mL), BIO (0 or 0.15 µM), SB431542 (0 or 2 µM), and VEGF (0 or 50 ng/mL). Minitab Statistical Software was used to determine 32 experimental conditions that maximized the difference between conditions. After 7 days in culture, CD34^+^-hiPSC-EP yield was measured using FACS. A total of 100,000 cells were measured per experimental condition. A list of experimental conditions and their CD34^+^-hiPSC-EP yield can be found in Supplemental Table 1.

### 2D Culture of CD34^+^-hiPSC-EPs

500,000 CD34^+^-hiPSC-EPs from the GF protocol were centrifuged twice and resuspended in Complete Endothelial Growth Media 2 (EGM-2, PromoCell, C-22011) supplemented with 50 ng/mL VEGF, 10 µM Y-27632, and 100 U/mL penicillin-streptomycin (ThermoFisher Scientific, 15140122), termed encapsulation media. They were then plated at a density of 17,500 cells/cm^2^ onto vitronectin-coated 48-well plates. During the 7-day culture period, media was replaced daily with EGM-2 supplemented with 50 ng/mL VEGF, termed endothelial culture media, or with Complete Smooth Muscle Growth Medium (SMGM, Lonza, CC-3182), supplemented with 50 ng/mL Recombinant Human Platelet-Derived Growth Factor-BB (PDGF, R&D Systems, 220-BB-050).

### Encapsulation of CD34^+^-hiPSC-EPs in Collagen/NorHA IPN Hydrogels

A total of 500,000 CD34^+^-hiPSC-EPs were centrifuged twice and resuspended in encapsulation media. Collagen/norbornene-modified hyaluronic acid (NorHA) interpenetrating polymer network hydrogels (IPNs) were synthesized as described previously^14^. In brief, RGD (GCGYGRGDSPG) and an enzymatically degradable (DEG) peptide crosslinker (KCGPQGIWGQCK) were reconstituted in encapsulation media to concentrations of 40 mg/mL and 20 mg/mL, respectively. RGD, DEG, and additional encapsulation media were added to lyophilized NorHA (0.51 norbornene functionalization) to a final concentration of 10 mg/mL NorHA, 2 mM RGD, and 1.98 mM DEG. This DEG concentration crosslinks 25% of the available norbornene groups present on the hyaluronic acid, and we have previously shown that the resulting IPN hydrogel supports robust vascular network formation^14^. The photoinitiator lithium phenyl-2,4,6-trimethylbenzoylphosphinate (LAP, Sigma-Aldrich, 900889-1G) was dissolved in encapsulation media to a final concentration of 0.5 wt%. RGD, DEG, and NorHA were synthesized and characterized as described previously^14^.

To encapsulate CD34^+^-hiPSC-EPs, 10x Medium M199 (ThermoFisher Scientific, 11150059), Type I Collagen from Rat Tail (Corning, 354249), LAP, and additional encapsulation media were mixed. The Collagen was then neutralized with 1 M Sodium Hydroxide (Sigma-Aldrich, 72068-100ML). The NorHA solution was then added to the Collagen solution and mixed well. Finally, the cell solution was added and 50 µL of the IPN hydrogel solution was pipetted into each well of a µ-Slide 18 Well Glass Bottom (Ibidi, 81817) and was allowed to solidify at 37°C at 5% CO_2_. The IPN hydrogels were then exposed to 365 nm UV light at 10 mW/cm^2^ for 50 seconds and then fed with 100 µL of encapsulation media. 24 hours after cell encapsulation, media was replaced with endothelial culture media, which was replaced daily for 7 days. The final IPN hydrogel contains 1 wt% NorHA and 0.25 wt% Collagen.

### Immunocytochemistry of Cells and Networks in IPN Hydrogels

The vessel-like networks generated within the hydrogels were visualized by staining with rhodamine-phalloidin (1:40, ThermoFisher Scientific, R415) following an immunocytochemistry protocol previously described by us^14-16^. For network quantification, Z-stacks were acquired at 13 µm intervals on a spinning disk confocal microscope (Zeiss Axio Observer Z1 with Yokogawa CSU-X1M). At least 4 regions of interest across the full depth of the hydrogel were acquired per hydrogel sample, and at least three hydrogels were analyzed per experimental condition. Additionally, to visualize cell secreted Collagen IV, cells were stained with mouse anti-human Collagen IV (1:1000, Invitrogen, 14-9871-82) and donkey anti-mouse Alexa Fluor 488 (1:200, abcam, ab150105). Nuclei were counterstained with DAPI. Cell-laden hydrogels were imaged using a Leica DMI8 laser scanning confocal fluorescence microscope. Z-stacks were acquired at 63x magnification at 0.5 µm intervals. At least 5 regions of interest, containing an isolated cluster of between 5 and 10 cells, were acquired for each hydrogel, and each experimental condition consisted of at least 5 hydrogels. Collagen IV deposition was quantified in ImageJ by measuring the average intensity of the Collagen IV signal within 2 µm of the cell surface for each slice of the z-stack.

### Immunocytochemistry of 2D cultured CD34^+^ hiPSC-EPs

GF-derived CD34^+^-hiPSC-EPs plated in 2D were cultured for 7 days and then fixed and stained following an immunocytochemistry previously described by us^14-16^. We stained using rabbit anti-human Calponin I (1:400, abcam, ab46794) and mouse anti-human VE-Cadherin (1:200, Santa Cruz Biotechnology, SC-9989). We then incubated with donkey anti-mouse Alexa Fluor 488 (1:200, abcam, ab150105) and donkey anti-rabbit Alexa Fluor 647 (1:200, Invitrogen, A150075). Nuclei were then counterstained with DAPI. Cells were imaged on a Leica DMi8 fluorescence microscope. At least 6 regions of interest were acquired per well, and 6 wells were imaged per experimental condition. We used ImageJ to quantify the proportion of endothelial (VE-Cadherin^+^) and smooth muscle (Calponin I^+^) cells among the total cell population in each region of interest.

### Length and Connectivity Analysis of Vessel-Like Networks

To analyze the total length, connectivity, and vessel diameter of the networks, we used a computational pipeline we previously developed^14^. Briefly, confocal z-stacks are filtered and binarized using ImageJ and then analyzed with a MATLAB script that converts the images into a nodal graph that describes the capillary-like structures as nodes (branch/endpoints) and links (vessels).

### Quantitative Reverse Transcription–Polymerase Chain Reaction

A total of 500,000 CD34^+^-hiPSC-EPs from both differentiation protocols as well as hiPSCs were centrifuged twice and messenger RNA (mRNA) was isolated using an RNeasy Mini Kit (Qiagen, 74106) and then reverse transcribed into complementary DNA with a High-Capacity cDNA Reverse Transcription Kit (ThermoFisher Scientific, 4368814) according to the manufacturers*’* instructions. Quantitative reverse transcription–polymerase chain reaction (qPCR) was performed with PowerUp SYBR Green (ThermoFisher Scientific, A25742) using the StepOne Plus system (Applied Biosystems), as described previously^15^. Relative expression of mRNA was quantified using the ΔΔCT method with GAPDH as the endogenous control. Results are presented as mean and standard deviation of three technical replicates. A list of primers used is given in Supplementary Table 2.

### 3*’* Tag-Seq

mRNA was collected from CD34^+^-hiPSC-EPs from both differentiation protocols immediately after FACS as described above. 3*’* Tag-Seq was performed by the University of Texas at Austin Genomics Sequencing and Analysis Facility (Center for Biomedical Research Support, RRID: SCR 021713) using an Illumina NovaSeq 6000 with 5 million reads per sample and 100 base pairs per read. The files were preprocessed by the Bioinformatics Consulting Group at the University of Texas at Austin and were mapped using the RNAseq pipeline by nf-core. The RNAseq pipeline aligned the reads to the GRCh38 human reference genome using *‘*STAR*’*, and gene counts were obtained using *‘*Salmon*’*.

Differentially expressed genes were determined using DESEQ2 with the adaptive shrinkage estimator, *‘*ashr*’*. Differentially expressed genes were defined as genes that had an adjusted p-value of 0.05 and a log fold change cut-off of 0.5. Using the differentially expressed genes, pre-ranked gene set enrichment analysis (GSEA) was used to determine enriched pathways using the Gene Ontology (GO) pathway set and a custom pathway using genes found to be involved in endothelial to mesenchymal transition (EndMT). Enriched pathways were determined by a false discovery rate (FDR) of 0.25. Differences in expression for individual genes are given using log_2_(fold change) of GF-derived cells relative to SM-derived cells, and differences in pathways are given using normalized enrichment scores (NES).

### Measuring Total MMP Activity

CD34^+^-hiPSC-EPs from both differentiation protocols were encapsulated in IPN hydrogels and cultured for a total of 7 days with daily media changes. Conditioned media from at least 6 hydrogels were collected for each condition every day. Total MMP activity was measured for each cell type each day using an MMP Activity Assay Kit (abcam, ab112146) using the manufacturer*’*s recommendation for end point analysis. Endothelial culture media was used as a negative control.

### Measuring Protease and Protease Inhibitor Secretion

To measure differences in release of proteases and protease inhibitors between CD34^+^-hiPSC-EPs from the two protocols, we used a Proteome Profiler Human Protease/Protease Inhibitor Assay Kit (R&D Systems, ARY025) using the recommended protocol for cell supernatant. Samples were prepared as follows: SM-derived hiPSC-EPs and GF-derived hiPSC-EPs were encapsulated in IPN hydrogels and conditioned media from at least 6 hydrogels per experimental condition were collected after 7 days of culture. Arrays were imaged using a MyECL Chemiluminescence Imager (ThermoFisher Scientific) with a 10-minute exposure time. Relative proteases and protease inhibitor levels were quantified using ImageJ. A full list of proteases and protease inhibitors measured using this array can be found in Supplementary Tables 3 and 4, respectively.

## Results

### Optimization of GF Differentiation

We first sought to determine optimal conditions for differentiating our chosen hiPSC line (DF19-9-11T; WiCell) into CD34^+^ cells using the GF protocol. In the paper that originally described the GF protocol, the researchers derived their own hiPSCs^1^. We hypothesized that differences in DNA methylation between different hiPSC lines would affect differentiation efficiency^17^; as a result, we first determined the most important factors for successful CD34^+^ cell differentiation.

We used Minitab statistical software to generate 32 experimental conditions to measure the effects of seven factors at two levels: seeding density, B27 supplement, Activin A, BMP4, BIO, SB431542, and VEGF. We chose two seeding densities, either 10,000 or 40,000 cells/cm^2^. Jalilian *et al* originally seeded hiPSCs at densities between 38,000 and 118,000 cells/cm^2^, with CD34^+^ cell yield increasing with seeding density. However, these seeding densities are all higher than the seeding density used in the SM protocol (20,000 cell/cm^2^), and we hypothesized that this discrepancy may be in part due to differences in proliferation rates between different hiPSC lines^18^. Because of this, we instead chose to test either half or double the seeding density used in the SM protocol. Because other groups have shown that Insulin withdrawal can improve hiPSC differentiation to hematopoietic stem cells^19^, we tested B27 with and without Insulin. All other factors were either not included or added at the concentration given in the original paper in order to identify any redundant factors in the protocol. For example, Activin A, BMP4, and BIO have all been used to induce mesoderm formation^20-24^, and we hypothesized that the differentiation could be successful using only one or two of these factors. We found that the two statistically significant factors for GF differentiation are Activin-A and VEGF (**Supplemental Figure 1**). In addition, the results showed that the optimal set of conditions for this differentiation protocol were as follows: seeding at 10,000 cells/cm^2^, using B27 minus Insulin, and using Activin A, BMP4, BIO, SB431542, and VEGF to differentiate the cells. This follows the original research apart from using B27 minus Insulin and decreasing the seeding density.

**Figure 1.**
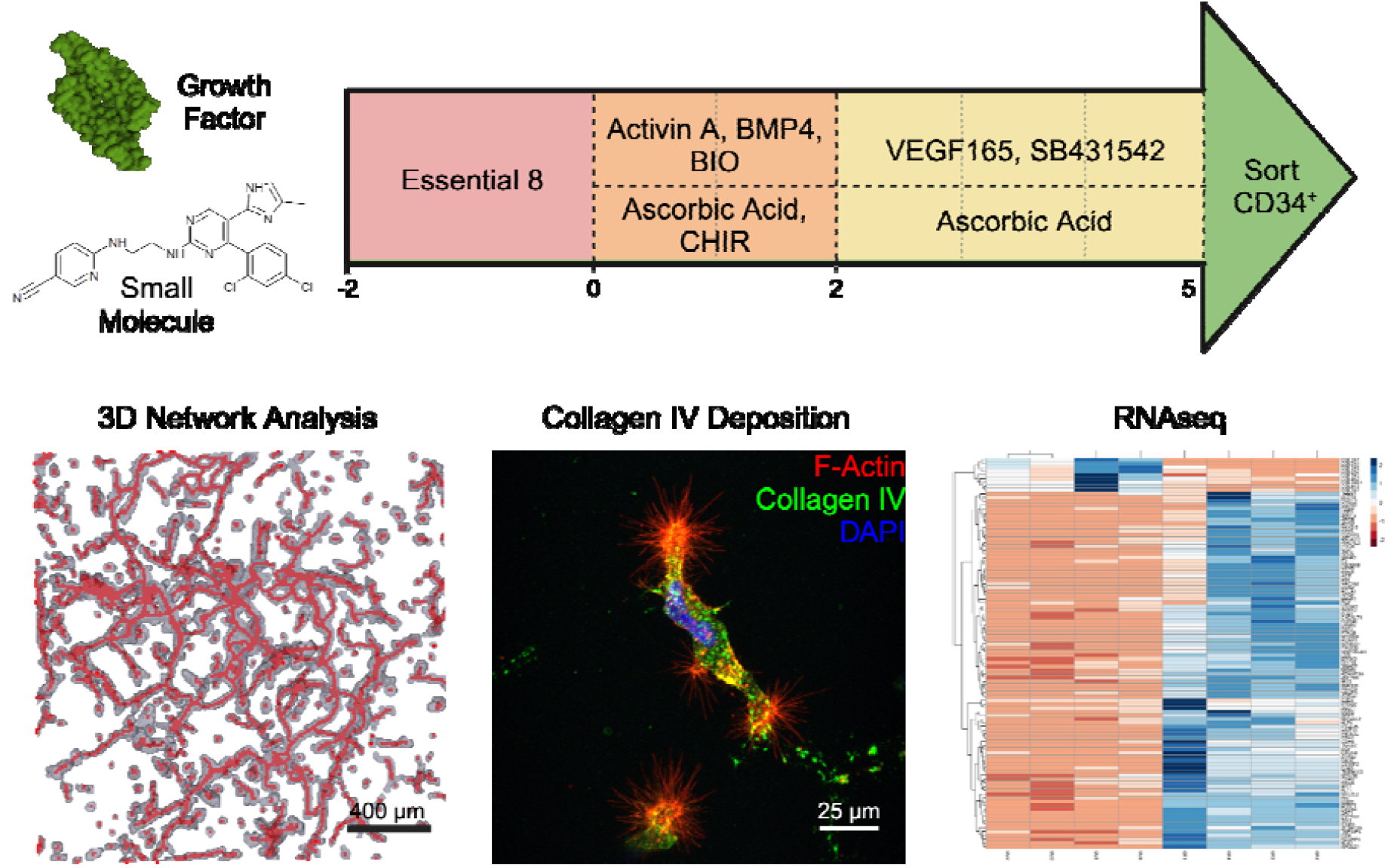
Experimental Overview: We generate CD34^+^-hiPSC-EPs using a differentiation protocol that primarily uses either small molecules or growth factors. CD34^+^-hiPSC-EPs are then encapsulated in Collagen I/Norbornene-modified hyaluronic acid hydrogels and compare the two cell types using our computational pipeline to evaluate vasculogenesis, RNAseq to compare gene expression, and Collagen deposition and protease activity to evaluate extracellular matrix remodeling. Created using BioRender.com.

### GF Protocol Yields Bipotent CD34^+^-hiPSC-EPs

To verify the multipotency of the GF derived cells, we plated CD34^+^-hiPSC-EPs onto vitronectin-coated plates and cultured the cells for 7 days in either EGM-2 or SMGM-2. Previously, we demonstrated that SM-derived CD34^+^-hiPSC-EPs can differentiate into both endothelial cells and smooth muscle cells by culturing in either EGM-2 or SMGM-2, respectively^15^. Similar to results in SM-derived cells, we found that GF-derived cells differentiated into both endothelial and smooth muscle cells regardless of culture media **(Supplemental Figure 2**). However, a higher percentage of cells cultured in EGM-2 expressed VE-Cadherin compared to cells cultured in SMGM-2 (77.1% vs 60.5%, p=0.0017).

**Figure 2.**
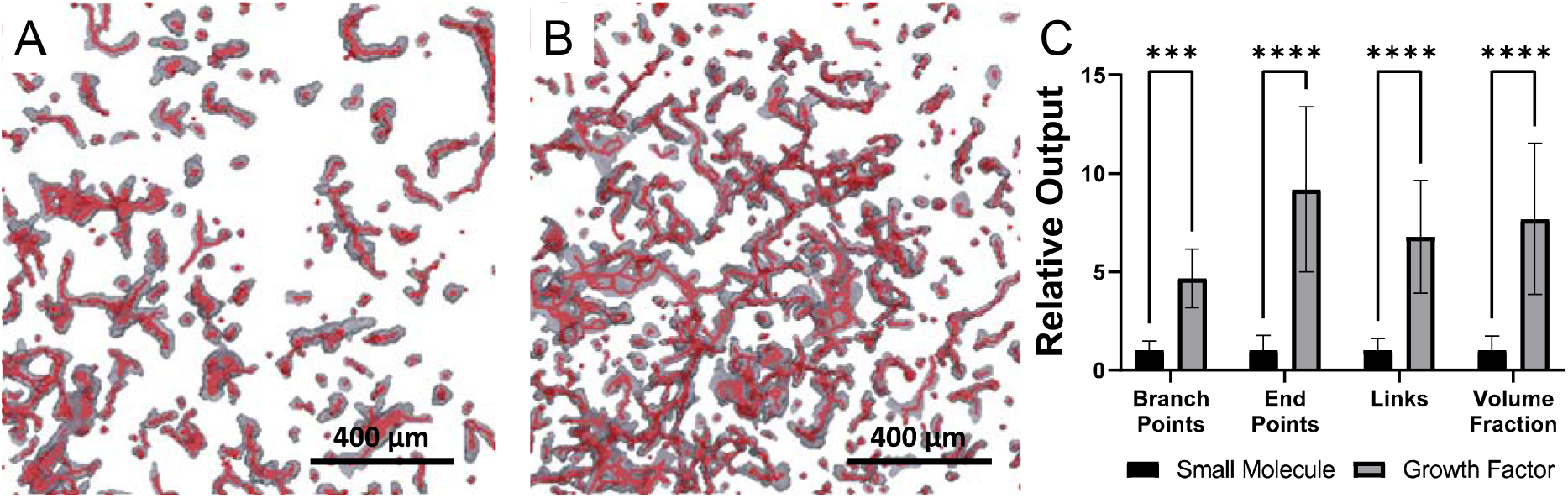
Growth Factor-Derived Cells have Higher Vasculogenic Potential: We generated CD34^+^-hiPSC-EPs using either the small molecule (A) or growth factor (B) protocol and encapsulated the cells in Collagen I/Norbornene-modified hyaluronic acid interpenetrating polymer network hydrogels (IPNs). After 7 days in culture, growth factor protocol-derived cells have significantly higher numbers of branch points, end points, links, and volume fraction containing vessel-forming cells (C). This indicates that growth factor-derived cells have a superior vasculogenic potential.

### GF-Derived CD34^+^-hiPSC-EPs Exhibit Higher Vasculogenic Potential

Next, we wanted to compare the vasculogenic potential of SM- and GF-derived CD34^+^-hiPSC-EPs. We encapsulated these cells in Collagen/NorHA interpenetrating network hydrogels (Coll/NorHA IPNs). We have previously demonstrated that SM-derived cells undergo vasculogenesis in both pure Collagen and Coll/NorHA IPNs^14-16,25^, but we chose to perform all experiments in Coll/NorHA IPNs in order to avoid hydrogel compaction that can occur in hydrogels containing only Collagen^14^.

Overall, we found that GF-derived CD34^+^-hiPSC-EPs formed more robust vasculature compared to SM CD34^+^-hiPSC-EPs (**Figure 2a-b**). Our computational analysis showed that, relative to SM-derived cell-laden hydrogels, GF-derived cell-laden hydrogels had 4.66 times as many branch points (p=0.0006), 9.19 times as many end points (p<0.0001), 6.78 times as many links (p<0.0001), and 7.68 times the volume of the hydrogel containing vessels (p<0.0001), as shown in **Figure 2c**. This demonstrates both that GF-derived CD34^+^-hiPSC-EPs are able to form vasculature in Coll/NorHA IPNs and that there are significant differences in the vasculogenic potential of the CD34^+^-hiPSC-EPs depending on their derivation protocol.

### SM- and GF-Derived CD34^+^-hiPSC-EPs Have Different Endothelial Gene Expression Profiles

To determine differences in gene expression between SM- and GF-derived CD34^+^-hiPSC-EPs, we extracted RNA from both cell types immediately after FACS and measured the expression of several endothelial cell-associated genes. In GF-derived cells, relative to SM-derived cells, we observed a 3.17-fold increase in *CD34* (p<0.0001), a 2.39-fold increase in *KDR* (p<0.0001), a 2.29-fold increase in *CDH5* (p<0.0001), a 1.97-fold increase in *PDGFRB* (p<0.0001), a 2.98-fold increase in *NOTCH1* (p<0.0001), a 6.97-fold increase in EFNB2 (p<0.0001), and a 3.11-fold increase in *EPHB4* (p<0.0001). There was no statistically significant difference in the expression of *CD31* and *NRP2*. Increased expression of the arterial markers *NOTCH1* and *EFNB2* as well as the venous marker *EPHB4* suggest that GF-derived cells are beginning to further differentiate into arterial and venous subtypes (**Supplemental Figure 3**). Overall, the increased expression of multiple endothelial genes indicates that GF-derived cells are more similar to mature endothelial cells; however, the increased expression of the endothelial progenitor marker *CD34*, suggests otherwise. We have previously shown a decrease in *CD34* expression in CD34^+^-hiPSC-EPs following encapsulation and culture in Collagen hydrogels as the cells mature, and Patel *et al* showed that this phenomenon also occurs *in vivo* Because of these contradictory trends, we then performed further gene expression analysis to investigate the phenotype of GF-derived cells.

**Figure 3.**
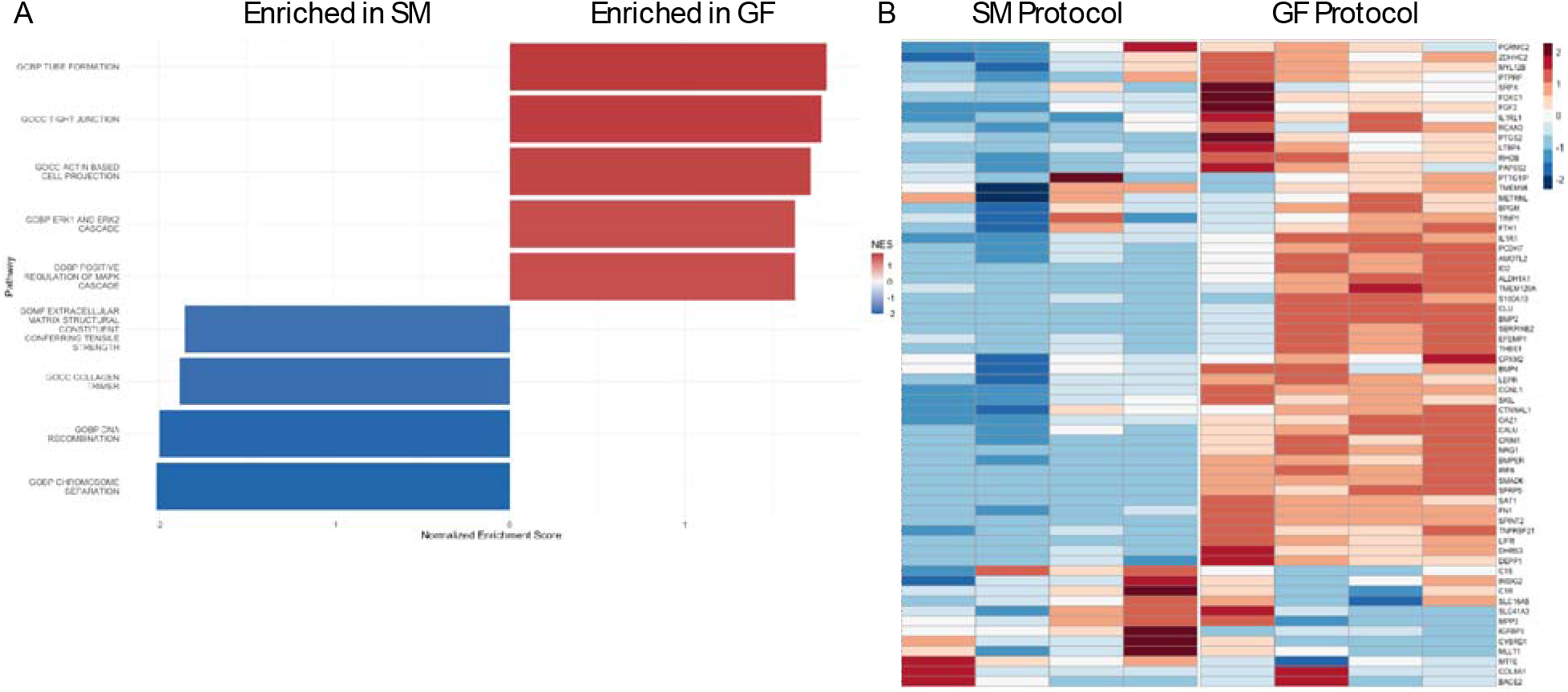
Overview of TagSeq Results: (A) GF-derived cells are enriched in pathways associated with cell migration (Actin-Based Cell Projections) and angiogenesis (Tube Formation, ERK1/ERK2 Cascade, MAPK Cascade, Tight Junctions), whereas SM-derived cells are enriched in pathways associated with ECM remodeling (Extracellular Matrix Structural Constituent Conferring Tensile Strength, Collagen Trimer) and cell proliferation (DNA Recombination, Chromosome Separation). (B) Heatmap of genes associated with endothelial-to-mesenchymal transitions shows that GF-derived cells are likely undergoing an endothelial-to-mesenchymal transition.

### GF-Derived CD34^+^ -hiPSC-EPs Endothelial-to-Mesenchymal Transition

To further explore transcript-level differences between the two cell types, we extracted RNA as described above and performed 3*’* Tag-Seq in cooperation with the University of Texas at Austin Genomics Sequencing and Analysis Facility. All data analysis was performed using an established analysis pipeline. We chose to use 3*’* Tag-Seq because it provides comparable accuracy to bulk RNA sequencing while requiring fewer reads, which reduces cost^26^. We focused our analysis on two main areas: arterial/venous specification and the endothelial-to-mesenchymal transition. We were interested in the former because our qPCR results suggest that the GF-derived cells are beginning to differentiate into arterial and venous endothelial cells. EndMT results in endothelial subtypes with increased *CD34* expression and cell migration^27^.

Based on our qPCR data, which showed higher expression of arterial genes compared to venous genes in GF-derived cells, we expected GF-derived CD34^+^-hiPSC-EPs to favor the arterial subtype. Because artery and vein formation both involve similar genes, we chose to exclude genes present in both arterial and venous pathways. We found that GF-derived cells are enriched in the pathway for arterial morphogenesis (NES=1.22, FDR=0.219). Some of the enriched genes include *LRP2*^28^, *FOLR1*^29^, *APOB*^30^, and *FOXH1*^31^, which are involved in formation of the aorta during embryonic development, and TGFB2. In addition, we found that SM-derived CD34^+^-hiPSC-EPs were enriched in pathways involved in the cell cycle, such as those for chromosome separation (M19628, NES =-2.02, FDR=0.094) and DNA recombination (GO:0006310, NES=-2.00, FDR=0.108) and this provides further evidence that GF-derived CD34^+^-hiPSC-EPs are more similar to arterial endothelial cells. Chavkin *et al* showed that a lower percentage of endothelial cells found in arteries were in the S/G2/M phases of the cell cycle, meaning they were non-proliferative^32^. However, it is not likely that SM-derived CD34^+^-hiPSC-EPs are favoring the venous subtype because we did not observe enrichment in any venous-specific markers. Taken together, this indicates that the GF protocol produces both arterial and venous endothelial cells and that the differences observed between cells of the two differentiation protocols are more likely due to the more stem-like phenotype of SM-derived CD34^+^-hiPSC-EPs

Our TagSeq data also suggests that GF-derived CD34^+^-hiPSC-EPs are favoring EndMT. EndMT is a process in which endothelial cells detach from their monolayer and adopt a mesenchymal phenotype^33^. Some initial evidence for this phenomenon is GF-derived CD34^+^-hiPSC-EPs*’* enrichment in pathways involved in cell migration and angiogenesis, such as actin-based cell projections (GO:0098858, NES=1.72, FDR=0.0767), cell-cell junction organization (GO:0034330, NES=1.7, FDR=0.0786), and tube formation (GO:0035148, NES=1.81, FDR=0.0356). In addition, cells undergoing EndMT are enriched in *CD34*, matching the qPCR data. To further investigate this, we used a gene set for EndMT genes from Slenders *et al* and found that GF-derived cells were enriched in this pathway (NES=1.73, FDR=0.011)^34^. Enriched genes include those encoding proteins that induce EndMT, such as *BMP4, FGF2*, and *SKIL*^*3*5^, all of which activate Smad5, as well as genes upregulated from Smad5 activation, such as *SKIL*^36^, *SERPINE1*^35^, and *SPINT2*^37^, all associated with increased cell migration and angiogenesis.

One commonly associated phenotype for cells undergoing EndMT is an increase in extracellular matrix (ECM) deposition^33^. Interestingly, we found that SM-derived CD34^+^-hiPSC-EPs were enriched in multiple pathways for Collagen synthesis and deposition, such as Collagen biosynthesis and modifying enzymes (M26999, NES=-1.82, FDR=0.222) and complex of Collagen trimers (GO:0098644, NES=-1.95, FDR=0.146).

### SM-Derived CD34^+^-hiPSC-EPs More Actively Remodel Extracellular Matrix

Although SM-derived CD34^+^-hiPSC-EPs were enriched in genes for both ECM deposition and degradation, this data was generated using cells immediately following FACS, and we were interested in seeing if this phenomenon persisted following encapsulation in Collagen/NorHA IPNs. To investigate this, we collected conditioned media from CD34^+^-hiPSC-EP-laden hydrogels from both differentiation protocols and used a Proteome Profiler Human Protease/Protease Inhibitor Array to measure relative concentrations of multiple proteases and protease inhibitors. Because secreted proteins may vary throughout the 7-day culture period, we first used a total MMP Activity Assay to determine the time point with the highest MMP activity for use in the array. As expected, both the GF- and SM-derived CD34^+^-hiPSC-EPs had the highest MMP activity on day 7, so this time point was used for future experiments (**Supplemental Figure 4**).

**Figure 4.**
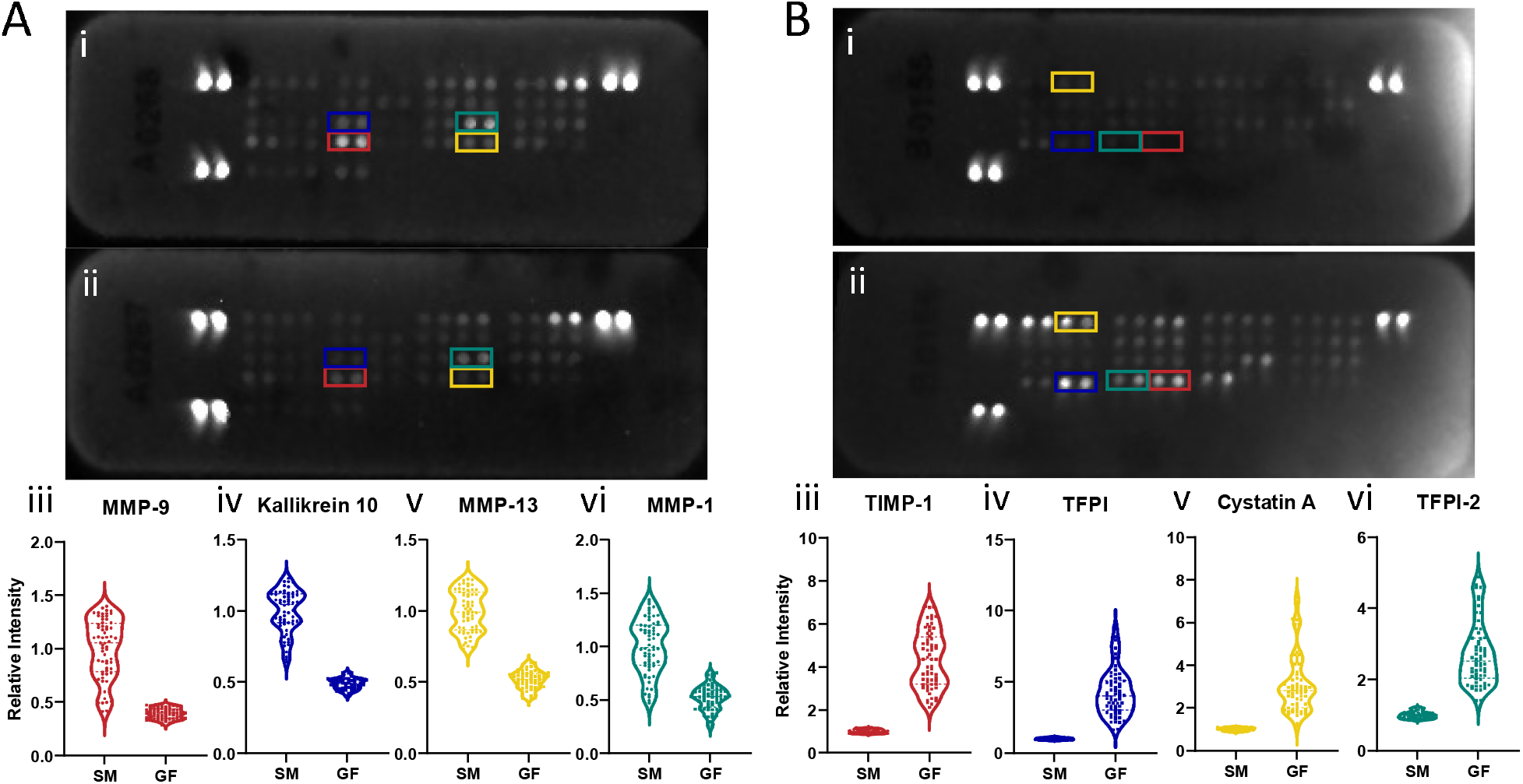
Increased Extracellular Matrix Degradation in Small Molecule-Derived Cells. CD34^+^-hiPSC-EPs from both differentiation protocols were encapsulated in Collagen I/Norbornene-modified hyaluronic acid hydrogels, and we collected conditioned media from at least 6 hydrogels per experimental condition to run two multiplexed arrays to measure proteases (A) and protease inhibitors (B). For both sets of arrays, (i) is the array for the small molecule protocol, and (ii) is the array for the growth factor protocol. In the protease array, we observed increased secretion of MMP-9 (iii, red), Kallekrein 10 (iv, blue), MMP-13 (v, yellow), and MMP-1 (vi, green) in small molecule-derived cells. In the protease inhibitor array, we observed increased secretion of TIMP-1 (iii, red), TFPI (iv, blue), Cystatin A (v, yellow), and TFPI-2 (vi, green) in growth factor-derived cells. This indicates increased extracellular matrix remodeling by small molecule-derived cells. Each dot in the violin plots represents the intensity of one pixel. n=2 for all conditions.

The protease array revealed substantial differences between the two differentiation protocols. Overall, SM-derived cells secrete higher concentrations of proteases compared to GF-derived cells. Specifically, SM-derived cells, compared to GF-derived cells, secreted higher levels of MMP-9 (2.54-fold), Kallikrein 10 (2.05-fold), MMP-13 (1.94-fold) and MMP-1 (1.92-fold) (**Figure 4a**). MMP-1 and MMP-13 both degrade Collagen I^38^, and we expected high concentrations of type-I Collagenases because our hydrogel system contains both Collagen I itself and a Collagenase-sensitive peptide crosslinker. MMP-9 degrades Collagen IV^38^, the most abundant form of Collagen in the endothelial basement membrane. Kallikrein 10 is expressed by endothelial cells and plays an anti-inflammatory role^39^. On the other hand, GF-derived cells had overall higher secretion of protease inhibitors. For example, GF-derived cells secreted 4.33 times more TIMP-1, 4.24 times more TFPI, 3.07 times more Cystatin A, and 2.71 times more TFPI-2 (**Figure 4b**). TIMP-1 inhibits Collagenases such as MMP-1, MMP-9, and MMP-13^40^. Cystatin A is a cysteine protease inhibitor that plays a role in ECM degradation during the initial stages of angiogenesis^41^. TFPI-1 and TFPI-2 are both inhibitors of the coagulation cascade; we believe that the cells are secreting these two proteins in response to heparin present in EGM-2^42^. Overall, this shows that SM-derived cells are more actively remodeling their ECM, matching the data from TagSeq.

#### SM-Derived CD34^+^ -hiPSC-EPs Secrete More Collagen IV

SM-derived cells*’* more active ECM degradation suggests an increase in ECM protein secretion. To test this as well as the finding from TagSeq that SM-derived cells are secreting more Collagen, we encapsulated CD34^+^-hiPSC-EPs from both differentiation protocols in Collagen/NorHA IPNs for 7 days, after which they were fixed and stained for F-Actin and Collagen IV. We located the cell boundaries using the F-Actin stain and measured the fluorescent intensity of the Collagen IV staining within 2 µm of the cell surface. We limited our measurements because other groups have shown that cells encapsulated for similar time frames deposit ECM within this distance^43,44^. As expected, we found that SM-derived cells secrete significantly more Collagen IV compared to GF-derived cells (753.5 vs 81.03, p<0.0001), as shown in **Figure 5**.

**Figure 5.**
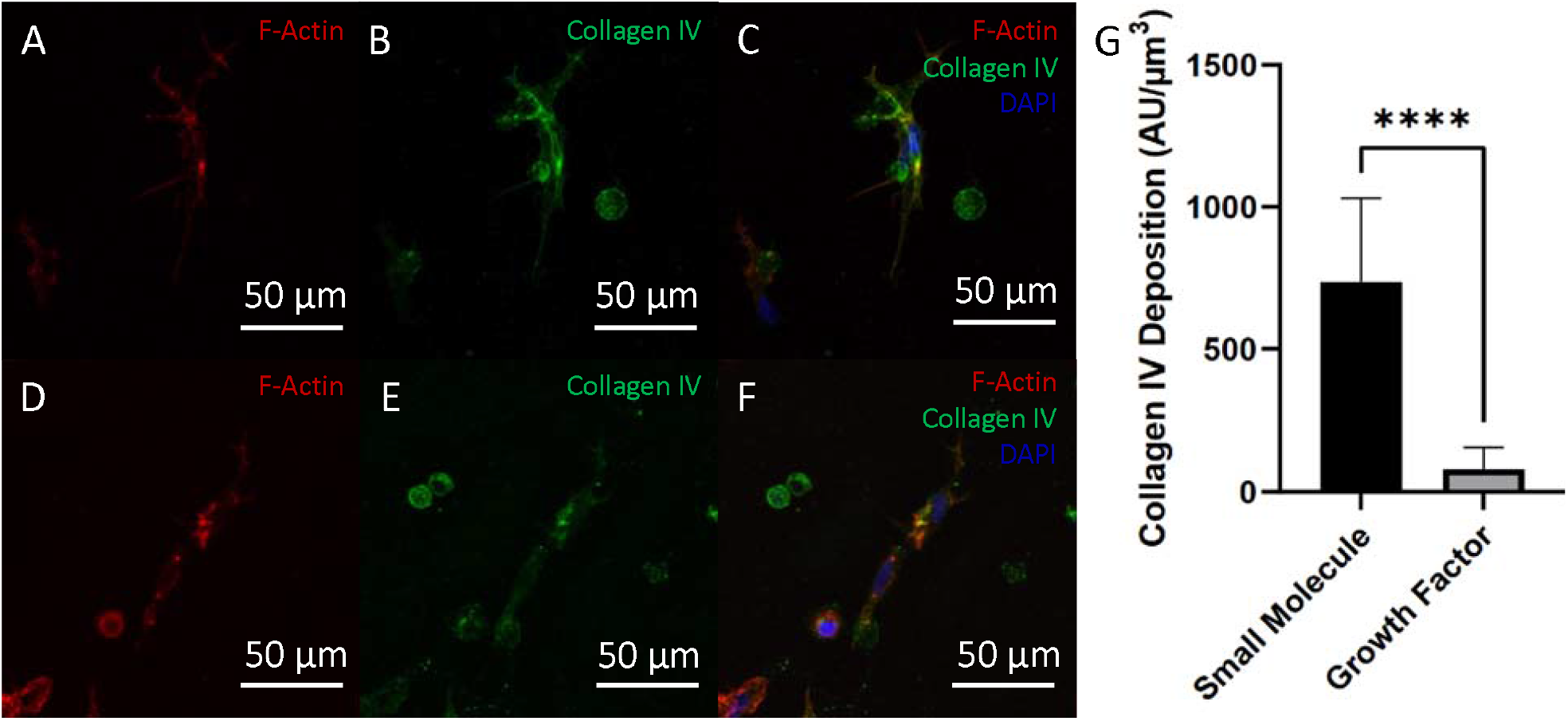
Small Molecule Derived Cells Deposit More Collagen IV. CD34^+^-hiPSC-EPs from both differentiation protocols were encapsulated in Collagen I/Norbornene-modified hyaluronic acid hydrogels, and we fixed and stained to visualize Collagen IV and F-Actin after 7 days in culture. We found that growth factor-derived cells secreted significantly more Collagen IV, which, combined with the protease/protease inhibitor array results, provides strong evidence of increased extracellular matrix remodeling in small molecule-derived cells.

## Discussion

These results demonstrate significant differences between two endothelial progenitor differentiation protocols at the gene, protein, and functional cell levels. Although multiple groups have created methods for self-described hiPSC-EP differentiation, there are significant differences between protocols regarding small molecules and growth factors administered, length of time for the differentiation (ranging from 5-25 days), and the choice of surface markers for identifying EPs, (such as CD34, CD31, CD144, and CD309^2-12,45^). In this study, we focused on isolating CD34^+^ cells using two different differentiation protocols: one that exclusively utilizes small molecules to drive differentiation, and another that utilizes a combination of growth factors and small molecules.

We initially performed qPCR on freshly isolated CD34^+^-hiPSC-EPs and encapsulated the cells in 3D Collagen/NorHA IPNs to investigate differences between the protocols; we found significant differences in vasculogenic potential (**Figure 2**) and gene expression (**Supplemental Figure 3**) that pointed to GF-derived cells*’* increased maturity, but we were unable to determine the exact phenotype of the GF-derived cells. However, it was clear that despite starting with the same hiPSC line and isolating EPs *using the same surface marker, the two protocols produced highly distinct cell populations*.

To further explore the specific cell types generated by these differentiation protocols produced, we performed gene expression analysis with 3*’*-TagSeq. Our initial findings supported the qPCR results, with enrichment in multiple pathways associated with angiogenesis in GF-derived CD34^+^-hiPSC-EPs. In addition, GF-derived CD34^+^-hiPSC-EPs were more similar to arterial endothelial cells. We expected to see some evidence of arterial/venous specification even at such an early time point, since in embryonic development endothelial cells commit to an arterial or venous fate even before the first vasculature forms^46^. In addition, other studies have used SB431542 and VEGF, two major components in the GF protocol, to drive differentiation to arterial endothelial cells^13^. Not surprisingly, GF-derived cells favored this endothelial subtype.

Interestingly, we found that GF-derived cells expressed genes associated with the endothelial-to-mesenchymal transition even though the cells retained tight junctions and formed lumenized vasculature when cultured in 2D and 3D, respectively, and neither of these features are associated with a mesenchymal fate^27^. However, EndMT does not always result in permanent commitment to a mesenchymal phenotype^47^; there are intermediate states in which cells only adopt some features of mesenchymal cells and later return to a quiescent endothelial phenotype. This is known as either partial-EndMT or endothelial-to-mesenchymal activation^47,48^. GF-derived cells retain expression of multiple endothelial-associated genes and form capillary-like vasculature, indicating that they are undergoing a partial EndMT. Two hallmarks for EndMT that were not observed in GF-derived cells were increased proliferation rates and increased ECM remodeling. The higher proliferation rate of SM-derived cells may be due to their immaturity relative to GF-derived cells; generally, as cells mature and become less stem-like, their proliferation rate decreases^49^.

Other reports have shown similar overall trends when investigating the effects of EndMT. For example, Park *et al* generated a partial knockout of Endoglin (Eng), a TGFβ receptor associated with EndMT, in mice and observed reduced angiogenesis, reduced cell migration, and increased Collagen IV deposition in Eng^+/-^ endothelial cells relative to those from wild-type mice^50^. Likewise, we have shown that SM-derived cells exhibit a phenotype similar to Eng^+/-^ endothelial cells relative to GF-derived cells, which are more similar to wild-type endothelial cells. The researchers found that TGFβ signaling activates Alk-5 in wild-type mice and Alk-1 in Eng^+/-^ mice, with the former having pro-angiogenic and anti-proliferative effects and the latter having the opposite effect. Although we did not observe significant differences in Endoglin expression in our TagSeq data, GF-derived cells are enriched in *TGFβ2*, which activates Endoglin, as well as *Smad5/6* and *SKIL*, both of which are downstream of Endoglin^51^. Taken together, this suggests that differences between SM- and GF-derived cells is at least partially attributable to canonical TGFβ signaling, resulting in a more activated phenotype similar to that seen *in vivo*

In conclusion, we generated CD34^+^-hiPSC-EPs from the same human induced pluripotent stem cell line using two differentiation protocols and found that the endothelial progenitors are highly distinct, ranging from the level of individual gene expression to their ability to form vasculature. SM-derived cells are less committed to becoming endothelial cells and more stem-like, whereas GF-derived cells are more similar to mature endothelial cells and are likely undergoing a partial endothelial-to-mesenchymal transition. This research highlights the benefits of developing different hiPSC-EP differentiation protocols in order to generate a wide variety of endothelial cell subtypes, such as arterial, venous, lymphatic, and tissue-specific endothelial cells. However, researchers that develop such protocols often do not investigate the target cell population beyond confirming their endothelial identity. There is a need for more thorough characterization of hiPSC-EP populations produced from different differentiation protocols to better understand the cell types present and to generate more physiologically accurate tissue constructs for *in vitro* models and future clinical applications.

## Author Contributions

BS and JZ contributed to study conception and design. Experiments were performed by BS and BL. Data analysis was performed by BS and SM. JZ and AB contributed resources, supervision, and project administration. The first draft of the manuscript was prepared by BS, and all authors contributed to revisions. All authors read and approved the final manuscript.

## Data Availability

3*’*-TagSeq data has been deposited in the Gene Expression Omnibus (GEO) under the accession code GSE278088. All other data that supports the findings of this study are available from the corresponding author upon reasonable request.

## Code Availability

Code used for processing 3*’*-TagSeq data can be accessed at https://nf-co.re/rnaseq/3.14.0/.

## Informed Consent

On behalf of all authors, the corresponding author states that informed consent was obtained from all participants involved in the study.

## Human and Animal Rights

On behalf of all authors, the corresponding author affirms that human and animal rights were upheld in the study.

## Acknowledgements

The authors thank Katie Hawlachs and Dr. Adrianne Rosales (University of Texas at Austin) for providing NorHA, RGD, and DEG, Dr. Jeanne Stachowiak (University of Texas at Austin) for use of her group*’*s spinning disk confocal microscope, and Dr. Cody Crosby (Southwestern University) for advice on optimizing the GF protocol. 3*’*-TagSeq was performed by the Genomic Sequencing and Analysis Facility at the University of Texas at Austin, Center for Biomedical Research Support (RRID: SCR_021713).

