## Supplemental Figure 1-4 for "Differentiation Protocol-Dependent Variability in hiPSC-Derived Endothelial Progenitor Functionality"

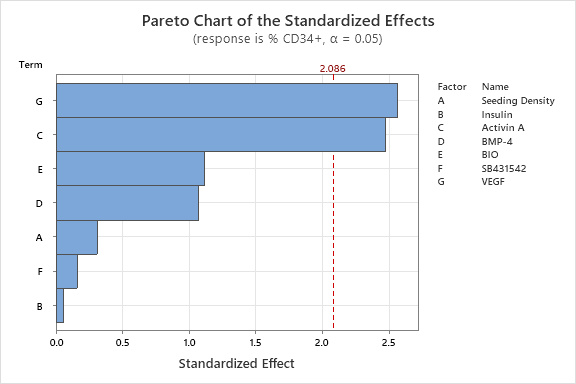

Supplemental Figure 1: Activin A and VEGF Significantly Effect CD34^+^ Yield in Growth Factor Protocol. Minitab Statistical Software was used to generate conditions and analyze results for a two-level factorial optimization experiment to determine the most important factors in the growth factor protocol. We found that only Activin A and VEGF had a statistically significant effect on CD34^+^ yield.

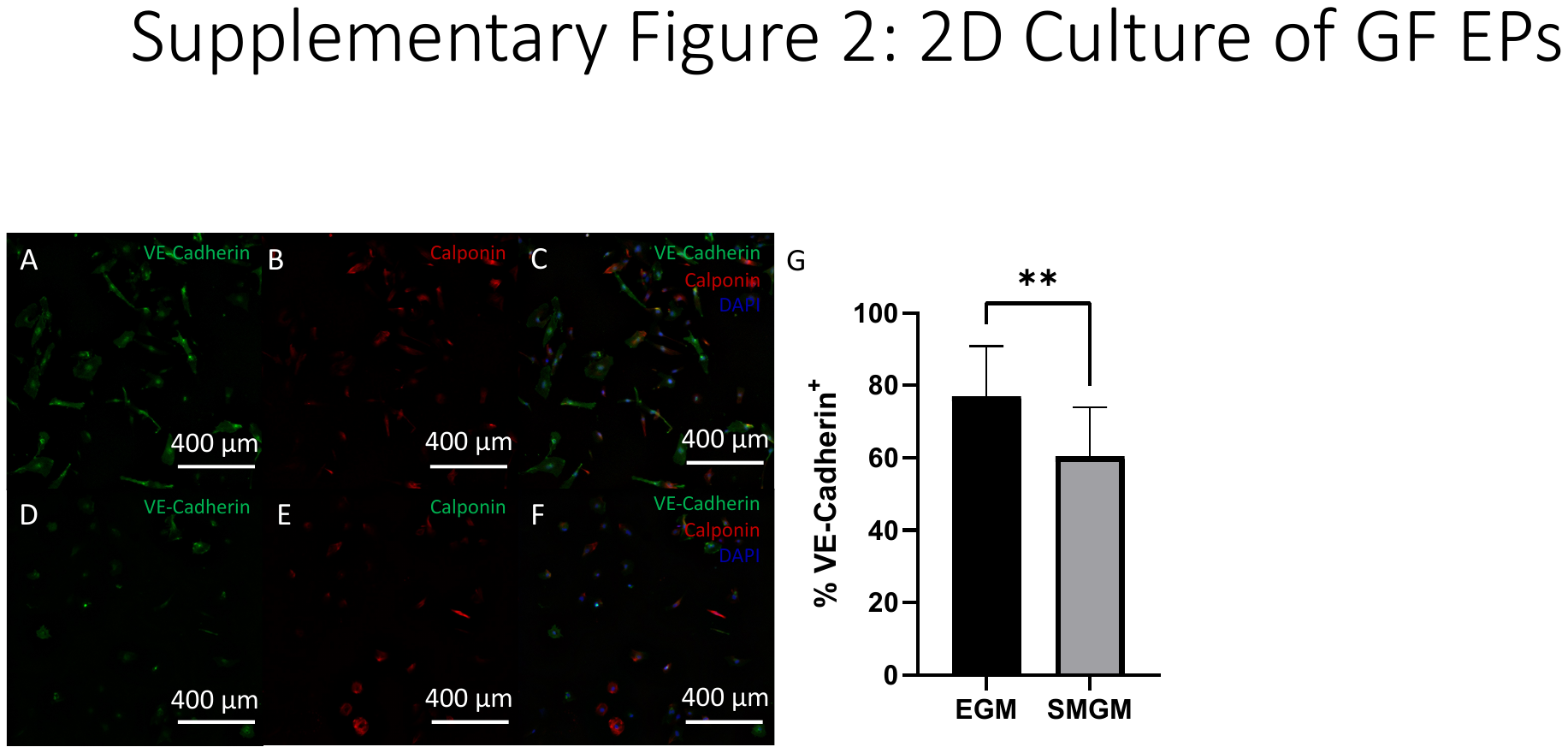

Supplemental Figure 2: Growth Factor-Derived CD34^+^-hiPSC-EPs are Bipotent. Growth factor-derived CD34^+^-hiPSC-EPs were seeded onto vitronectin-coated plates and cultured for 4 days with either endothelial growth media (A) (EGM) or smooth muscle growth media (B) (SMGM) to promote differentiation into either endothelial cells or smooth muscle cells, respectively. We then stained for VE-Cadherin to visualize endothelial cells and calponin to visualize smooth muscle cells and quantified the percentage of VE-Cadherin positive cells (C). We observed that while both media types supported differentiation into both endothelial and smooth muscle cells, there was a significantly higher percentage of VE-Cadherin positive cells in wells that received EGM. This demonstrates that growth-factor derived EPs are bipotent and that we can influence their differentiation through media supplementation.

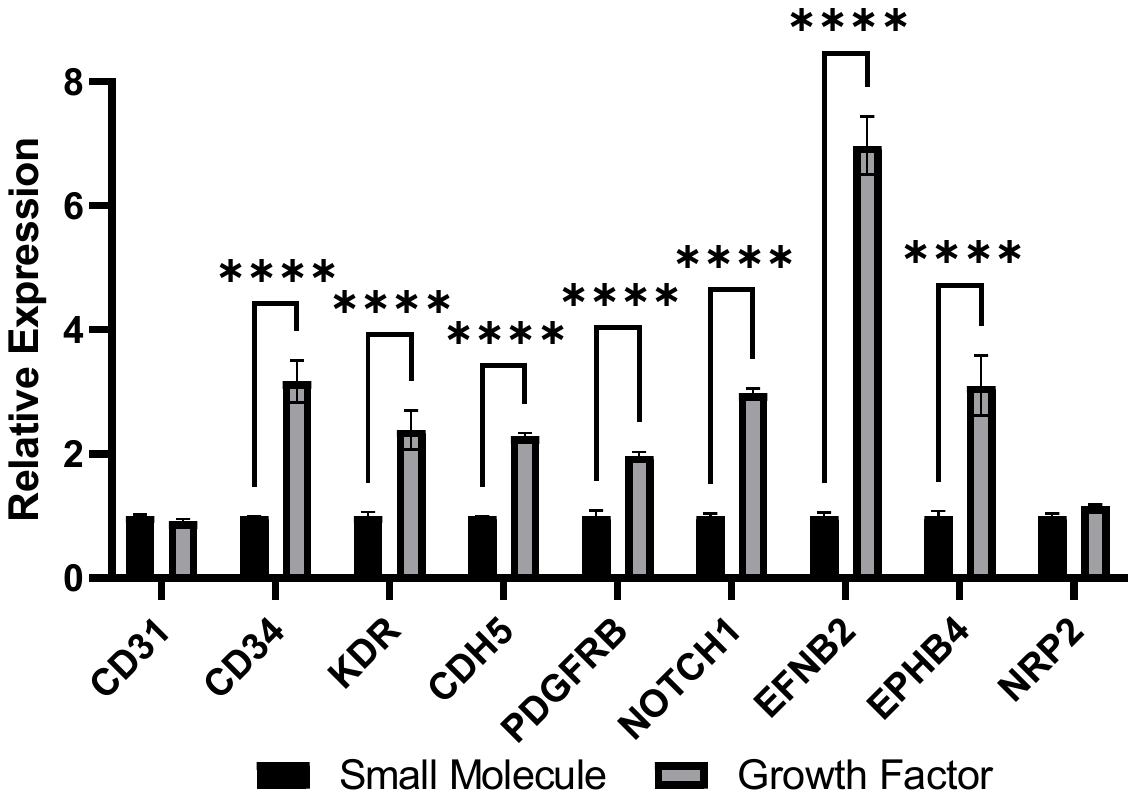

Supplemental Figure 3: Increased Maturity of GF-Derived CD34^+^-hiPSC-EPs. We extracted RNA from CD34^+^-hiPSC-EPs immediately after sorting and used qPCR to compare expression of several endothelial-related genes. We observed that growth factor-derived cells had higher expression of *CD34*, *CD31*, *KDR*, *CDH5*, *PDGFRB*, *NOTCH1*, *EFNB2*, and *EPHB4*. Higher expression of the first 5 listed genes suggests that the growth factor-derived cells are more similar to mature endothelial cells, and higher expression of the latter 3 suggests that the cells have started differentiating into arterial and venous subtypes.

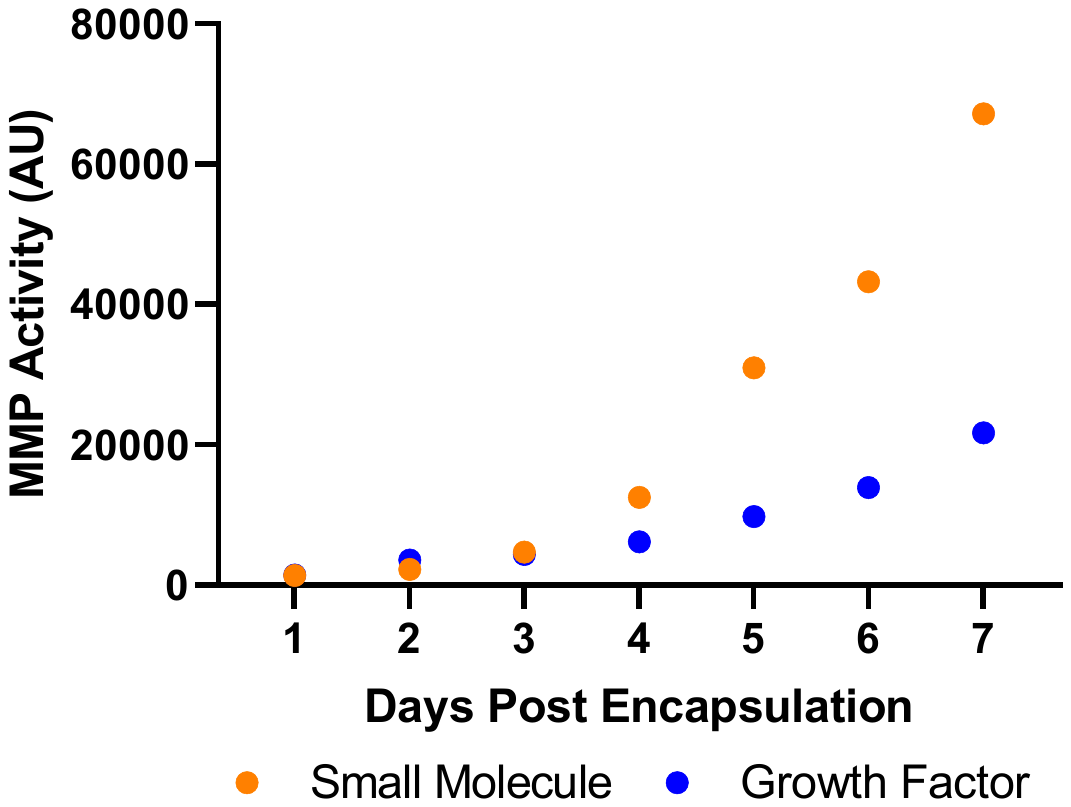

Supplemental Figure 4: Small Molecule-Derived Cells Have Higher Total MMP Activity: We encapsulated CD34^+^-hiPSC-EPs from both differentiation protocols and collected conditioned media daily for 7 days. We then used a total MMP Activity Assay Kit to measure total MMP activity to determine the optimal time point for the protease/protease inhibitor array. For both protocols, we found that MMP activity was similar for the first 3 days in culture. After that, small molecule-derived cells had greatly increased MMP activity compared to growth factor-derived cells, and by day 7, the small molecule condition had roughly 3 times higher MMP activity that the growth factor condition. Because MMP activity was at its highest for both protocols at day 7, we chose this time point for the protease/protease inhibitor array.
