## Supplemental Tables 1-5 for "Differentiation Protocol-Dependent Variability in hiPSC-Derived Endothelial Progenitor Functionality"

Supplemental Table 1: Complete List of Experimental Conditions for Factorial Optimization Experiment and CD34^+^ Cell Yield.

| **Seeding Density (cell/cm^2^)** | **B27 (Plus or Minus Insulin)** | **Activin A (ng/mL)** | **BMP4 (ng/mL)** | **BIO (µM)** | **SB431542 (µM)** | **VEGF (ng/mL)** | **% CD34^+^** |
| --- | --- | --- | --- | --- | --- | --- | --- |
| 10,000 | Minus | 0 | 0 | 0 | 0 | 0 | 0 |
| 10,000 | Minus | 0 | 0 | 0.15 | 2 | 50 | 0 |
| 10,000 | Minus | 0 | 30 | 0 | 2 | 50 | 0 |
| 10,000 | Minus | 0 | 30 | 0.15 | 0 | 50 | 0.74 |
| 10,000 | Minus | 25 | 0 | 0 | 2 | 0 | 0.16 |
| 10,000 | Minus | 25 | 0 | 0.15 | 0 | 0 | 0.3 |
| 10,000 | Plus | 25 | 0 | 0.15 | 2 | 50 | 9.8 |
| 10,000 | Minus | 25 | 30 | 0 | 0 | 50 | 1.39 |
| 10,000 | Plus | 0 | 0 | 0 | 2 | 0 | 0.08 |
| 10,000 | Plus | 0 | 0 | 0.15 | 0 | 50 | 0 |
| 10,000 | Plus | 0 | 30 | 0 | 0 | 0 | 0 |
| 10,000 | Plus | 25 | 30 | 0 | 2 | 0 | 0.3 |
| 10,000 | Plus | 25 | 30 | 0.15 | 2 | 0 | 0.01 |
| 10,000 | Plus | 25 | 30 | 0.15 | 0 | 50 | 7.5 |
| 40,000 | Minus | 0 | 0 | 0 | 2 | 50 | 0.37 |
| 40,000 | Minus | 0 | 0 | 0.15 | 0 | 0 | 0.01 |
| 40,000 | Minus | 0 | 30 | 0 | 0 | 0 | 0.05 |
| 40,000 | Minus | 25 | 0 | 0.15 | 2 | 0 | 0.01 |
| 40,000 | Minus | 25 | 30 | 0 | 2 | 50 | 8.27 |
| 40,000 | Minus | 25 | 30 | 0.15 | 0 | 50 | 6.88 |
| 40,000 | Minus | 25 | 30 | 0.15 | 2 | 0 | 0.13 |
| 40,000 | Plus | 0 | 0 | 0.15 | 2 | 50 | 0.03 |
| 40,000 | Plus | 0 | 30 | 0 | 2 | 50 | 0.11 |
| 40,000 | Plus | 0 | 30 | 0.15 | 2 | 0 | 0 |
| 40,000 | Plus | 0 | 30 | 0.15 | 0 | 0 | 0.01 |
| 40,000 | Plus | 25 | 0 | 0 | 0 | 0 | 0.01 |
| 40,000 | Plus | 25 | 0 | 0 | 0 | 50 | 0 |
| 40,000 | Plus | 25 | 0 | 0 | 0 | 50 | 0 |

Supplementary Table 2: List of Primers Used for qPCR:

| Gene | Sequence (F=Forward, R=Reverse) |
| --- | --- |
| *GAPDH* | F: 5’-GTC AGT GGT GGA CCT GAC CT-3’  R: 5’-CCC TGT TGC TGT AGC CAA AT-3’ |
| *CD31* | F: 5’-GCT GAC CCT TCT GCT CTG TT-3’  R: 5’-TGA GAG GTG GTG CTG ACA TC-3’ |
| *CD34* | F: 5’-CCT AAG TGA CAT CAA GGC AGA A-3’  R: 5’-GCA AGG AGC AGG GAG CAT A-3’ |
| *KDR* | F: 5’-GTG ACC AAC ATG GAG TCG TG-3’  R: 5’-TGC TTC ACA GAA GAC CAT GC-3’ |
| *CDH5* | F: 5’-AAG CGT GAG TCG CAA GAA TG-3’  R: 5’-TCT CCA GGT TTT CGC CAG TG-3’ |
| *PDGFRB* | F: 5’-TGG CAG AAG AAG CCA CGT T-3’  R: 5’-GGC CGT CAG AGC TCA CAG A-3’ |
| *NOTCH1* | F: 5’-CAC GCG GAT TAA TTT GCA TCT G-3’  R: 5’-TCT TGG CAT ACA CAC TCC GAG AAC-3’ |
| *EPFNB2* | F: 5’-CCC AGT GAC ATT ATC ATC CC-3’  R: 5’-CAT CTC CTG GAC GAT GTA CAC C-3’ |
| *EPHB4* | F: 5’-AGA GGC CGT ACT GGG ACA TGA G-3’  R: 5’-TCC AGC ATG AGC TGG TGG AG-3’ |
| *NRP2* | F: 5’-GCA TGG CAA AAA CCA CAA GGT AT-3’  R: 5’-TGG AGC GTG GAG CTT GTT CA-3’ |

Supplementary Figure 3: List of Proteases Measured using Protease/Protease Inhibitor Array:

| ADAM8 | Cathepsin X/Z/P | MMP-7 |
| --- | --- | --- |
| ADAM9 | DPPIV/CD26 | MMP-8 |
| ADAMTS1 | Kallikrein 3/PSA | MMP-9 |
| ADAMTS13 | Kallikrein 5 | MMP-10 |
| Cathepsin A | Kallikrein 6 | MMP-12 |
| Cathepsin B | Kallikrein 7 | MMP-13 |
| Cathepsin C | Kallikrein 10 | Neprilysin/CD10 |
| Cathepsin D | Kallikrein 11 | Presenilin |
| Cathepsin E | Kallikrein 13 | Proprotein Convertase 9 |
| Cathepsin L | MMP-1 | Proetinase 3 |
| Cathepsin S | MMP-2 | uPA/Urokinase |
| Cathepsin V | MMP-3 |  |

Supplementary Table 5: List of Protease Inhibitors Measured using Protease/Protease Inhibitor Array:

| APP/Protease Nexin II (pan) | Lipocalin-1 | Serpin F1/PEDF |
| --- | --- | --- |
| Cystatin A | Lipocalin-2/NGAL | Testican 1/SPOCK1 |
| Cystatin B | RECK | Testican 2/SPOCK2 |
| Cystatin C | Serpin A5/Protein C Inhibitor | TFPI |
| Cystatin E/M | Serpin A8/Angiogensinogen | TFPI-2 |
| EMMPRIN/CD147 | Serpin A9/Centerin | TIMP-1 |
| Fetuin B | Serpin A12 | TIMP-2 |
| HAI-1 | Serpin B5/Maspin | TIMP-3 |
| HAI-2 | Serpin B6 | TIMP-4 |
| HE4/WFDC2 | Serpin B8/Proteinase Inhibitor 8 | Trappin-2/Elafin |
| Latexin | Serpin E1/PAI-1 |  |
